# Abolished frameshifting for predicted structure-stabilizing SARS-CoV-2 mutants: Implications to alternative conformations and their statistical structural analyses

**DOI:** 10.1101/2024.03.28.586935

**Authors:** Abhishek Dey, Shuting Yan, Tamar Schlick, Alain Laederach

## Abstract

The SARS-CoV-2 frameshifting element (FSE) has been intensely studied and explored as a therapeutic target for coronavirus diseases including COVID-19. Besides the intriguing virology, this small RNA is known to adopt many length-dependent conformations, as verified by multiple experimental and computational approaches. However, the role these alternative conformations play in the frameshifting mechanism and how to quantify this structural abundance has been an ongoing challenge. Here, we show by DMS and dual-luciferase functional assays that previously predicted FSE mutants (using the RAG graph theory approach) suppress structural transitions and abolish frameshifting. Furthermore, correlated mutation analysis of DMS data by three programs (DREEM, DRACO, and DANCE-MaP) reveals important differences in their estimation of specific RNA conformations, suggesting caution in the interpretation of such complex conformational landscapes. Overall, the abolished frameshifting in three different mutants confirms that all alternative conformations play a role in the pathways of ribosomal transition.

## Introduction

Programmed ribosomal frameshifting (PRF) is utilized by multiple RNA viruses to translate proteins from overlapping open reading frames (ORF) of their genome (Brierley 1995; Dinman 2006). This mechanism involves repositioning of the translating ribosome by one or two nucleotides, thus effectively changing the frame of translation (Atkins et al. 2016; Staple and Butcher 2003). One of the most common types of programmed ribosomal frameshifting is -1 PRF where the ribosome is forced to move back by 1 nucleotide which further continues protein synthesis by reading ORF in -1 frame (Dinman 2012). This kind of regulation enables the virus to encode multiple proteins (Bhatt et al. 2021). Severe Acute Respiratory Syndrome Coronavirus-2 (SARS-CoV-2) employs this -1 PRF to regulate the translation of proteins from its alternative overlapping open reading frames ORF1a and ORF 1b (Bhatt et al. 2021; K. Zhang et al. 2021; Kelly et al. 2020; Schlick, Zhu, Dey, et al. 2021).

The genetic makeup of SARS-CoV-2 is composed of single stranded positive sense RNA. Out of its 10 open reading frames, partially overlapped ORF1a and ORF1b of SARS-CoV-2 code for non-structural proteins including RNA-dependent RNA polymerase (RdRP), an essential enzyme critical for maintaining the viral life cycle inside the host cell (Subissi et al. 2014; Malone et al. 2022).

Translation of RdRP from ORF1b is facilitated by the -1 PRF which is triggered by a specific mRNA structure present at the interface of ORF1a and ORF1b (Subissi et al. 2014). Additionally, a hepta-nucleotide sequence known as "slippery site” is joined to this RNA structural element at its 5’ end by a short spacer region thus constituting the RNA frameshift signaling element (FSE) which regulates the -1 PRF in SARS-CoV-2 (Kelly et al. 2020). As mentioned above, pausing during ribosomal translation is followed by subsequent unfolding in the downstream RNA structure to continue protein translation from a new ORF.

Previous biophysical studies have determined that the three-stem H-type pseudoknot is the most likely RNA structure present in the FSE of SARS-CoV-2 RNA genome (Bhatt et al. 2021; K. Zhang et al. 2021; Jones and Ferré-D’Amaré 2022; Roman et al. 2021). A 6.9 Å cryo-EM structure of 88nt long SARS-CoV-2 FSE forms a δ shaped conformation with the 5’ end being flexible enough to switch between “threaded” and “unthreaded” conformation, thus regulating the FSE activity (K. Zhang et al. 2021). Two recent crystal structures of SARS-CoV-2 FSE also identified it as a H-type pseudoknot; however, these RNA FSE include deletions of certain residues to stabilize their crystal structures (PDB ID: 7MLX and 7LYJ) (Roman et al. 2021; 2022). Our earlier study which includes the combination of graph-based modelling, two-dimensional structure prediction algorithms, and chemical probing and mutational profiling studies on multiple FSE lengths of SARS-CoV-2 highlighted the presence of at least three conformations that SARS-CoV-2 FSE can adopt (Schlick, Zhu, Dey, et al. 2021). Significantly, we identified three alternative RNA FSE structures in our prior study of 77 nucleotide (nt) RNA constructs (Schlick, Zhu, Dey, et al. 2021). In addition to two different H-type pseudoknots (3_6 and 3_3 dual graphs), an unknotted 3-way junction (3_5 graph) was also identified in the FSE landscape (Schlick, Zhu, Dey, et al. 2021).

The extensive body of existing literature on the SARS-CoV-2 FSE makes it an ideal model system for understanding the role of multiple conformations in RNA processes. Here we study multiple FSE constructs, including structure stabilizing mutants predicted earlier in (Schlick, Zhu, Dey, et al. 2021), by DMS probing coupled with mutational profiling data and corresponding frameshifting efficiency assays. The abolished frameshifting on these mutants and consensus analysis using several algorithms that deconvolute the DMS chemical reactivity into multiple conformations (including, DANCE-MaP (Olson et al. 2022), DREEM (Tomezsko et al. 2020) and DRACO (Morandi et al. 2021)) underscores the importance of conformational transitions during frameshifting and the potential for new therapeutic strategies that can be developed based on selected RNA fold distributions.

## Results

### DMS FSE data are consistent with alternative secondary structures

We begin our investigation of the SARS-CoV-2 FSE secondary structure landscape by evaluating solved three-dimensional structures. In Figure 1A we show the structural model based on a cryo-EM of 87nt SARS-CoV-2 FSE construct including the slippery sequence (PDB ID 6XRZ) (K. Zhang et al. 2021).

**Figure 1:**
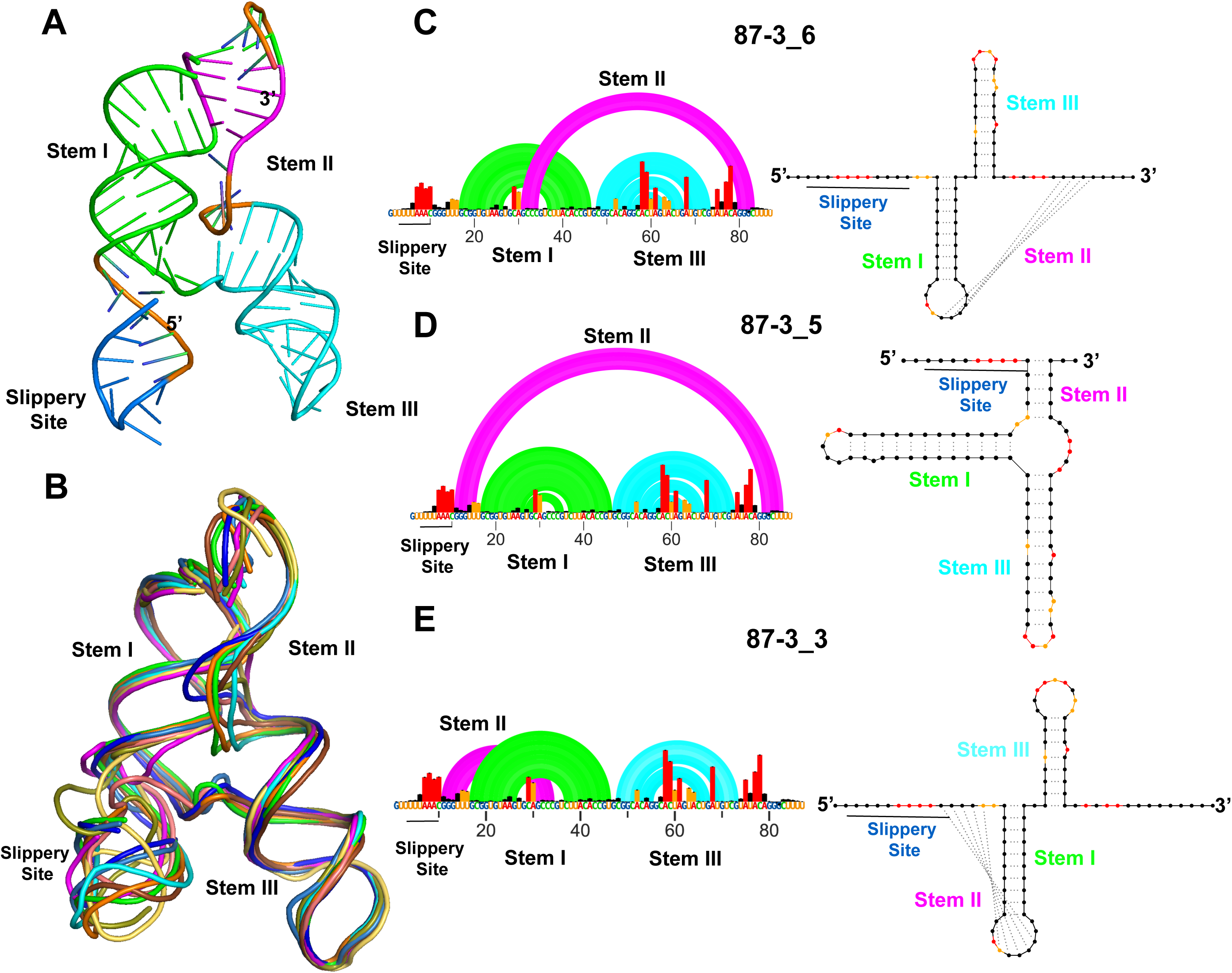
Conformational landscape of 87nt SARS-CoV-2 FSE. (A) 3D structure displaying 3_6 H-type pseudoknot conformation in λ form obtained through cryo-EM (PDB ID 6XRZ) (B) Multiple cryo-EM derived structural models displaying conformational consistency in 87nt FSE. The 5’ end consisting of slippery site is highly flexible. Arc plot and radial layout computed through DMS-MaP and SHAPEknots for 87nt FSE in (C) 3_6 H-type pseudoknot (D) 3_5 three-way junction and (E) 3_3 H-type pseudoknot. Slippery site, Stem I, Stem II, and Stem III are annotated in the arc plot and radial layout.

As can be seen in Figure 1B, several structural models were able to fit the cryo-EM density. There is high level structural agreement in the FSE region, with slightly more flexibility in the slippery site (Fig. 1A, 1B). Previous molecular dynamics simulations of this construct agree with this level of conformational dynamics, especially in stem helix 2 (Rangan et al. 2021; Schlick, Zhu, Dey, et al. 2021). In all these models however, the consensus secondary structure is the three stem 3_6 pseudoknot (H-type pseudoknot), Figure 1C.

The 3_6 notation refers to the RNA-as-graphs (RAG) notation of secondary structures, where dual graphs are indexed according to stem number at decreasing compactness, as introduced in (Fera et al. 2004; Jain, Tao, and Schlick 2020; Schlick, Zhu, Dey, et al. 2021). Our DMS-MaP (Dimethyl Sulfate-Mutational Profiling) experiments on this 87nt construct in Figure 1C indicate low (black), medium (yellow), and high (red) nucleotides based on their reactivity against DMS (Mustoe et al. 2019). We previously collected SHAPE-MaP (Selective 2’ hydroxyl acylation followed by Primer Extension-Mutational Profiling) data on this same construct (Schlick, Zhu, Dey, et al. 2021). Both DMS and SHAPE reactivities can be used as a pseudo-free energy term in thermodynamic folding algorithms to significantly improve structure prediction (Deigan et al. 2009; Greenwood and Heitsch 2020)

Our two DMS-MaP biological replicates in supplementary Figure S1 are quantitatively reproducible (R^2^>0.85). When used for structure prediction with SHAPEknots (Hajdin et al. 2013), we obtained three conformations, shown in Figures 1C, 1D and 1E. They correspond to 3_6, 3_5 and 3_3 RAG topologies, and their relative folding energies are within 3 kcal/mol (see Table 1). This result is remarkably consistent with our previous SHAPE-MaP data on 77nt, 87nt, and 144nt constructs, suggesting that the SARS-CoV-2 FSE adopts multiple conformations (Schlick, Zhu, Dey, et al. 2021). These data are also consistent with other groups’ analyses of the FSE secondary structure (Lan et al. 2022; Pekarek et al. 2023; Yan et al. 2022; Rangan et al. 2021; Schlick, Zhu, Dey, et al. 2021). We have emphasized in our prior work that not only can the RNA fold into different conformations for the same length (Schlick, Zhu, Dey, et al. 2021; Yan et al. 2023; Schlick, Zhu, Jain, et al. 2021); the fold distribution is also highly sensitive to length, when residues are either added or deleted (Yan et al. 2023) .

**Table 1:** Minimum free energies computed for shorter SARS-CoV-2 FSE constructs after DMS-MaP and ShapeKnots.

| SHAPEKnots | 3_6 | 3_5 | 3_3 | 3_1 | 2_1 | 3_5* |
| --- | --- | --- | --- | --- | --- | --- |
| 87 (6XRZ) | -42.4 kcal/mol | -40.4 kcal/mol | -39.4 kcal/mol | - | - | - |
| 7LYJ (66 nt) | -45.0 kcal/mol | - | - | - | - | - |
| 7MLX (65 nt) | -32.9 kcal/mol | - | - | -30.3 kcal/mol | - | - |
| 77 | -42.5 kcal/mol | -40.8 kcal/mol | -40.2 kcal/mol | - | - | - |
| 3_6 Stabilizing | -32.8 kcal/mol | - | - | - | - | - |
| 3_5 Stabilizing | - | -39.9 kcal/mol | - | - | - | - |
| 3_3 Stabilizing | - | - | -24.3 kcal/mol | - | - | - |

### DMS data of FSE crystallization constructs

We further collected DMS data on crystal constructs (PDB ID 7MLX and 7LYJ) [12,13] . As can be seen in Fig. 2A, the 66nt construct of 7LYJ adopts the 3_6 pseudoknot. Our DMS data, when used as a pseudo-free energy restraint term, also leads to the same single low energy structure (Fig. 2B and Table 1) in 3_6 topology. The 65nt construct (PDB ID 7MLX) (Figure 2C), is also predicted to adopt the 3_6 conformation (Fig. 2D). However, DMS data also point to an alternative 3_1, higher in energy conformation (Table 1). These data and analyses confirm the dominance of the 3_6 pseudoknot for short FSE lengths which does not contain any slippery site.

**Figure 2:**
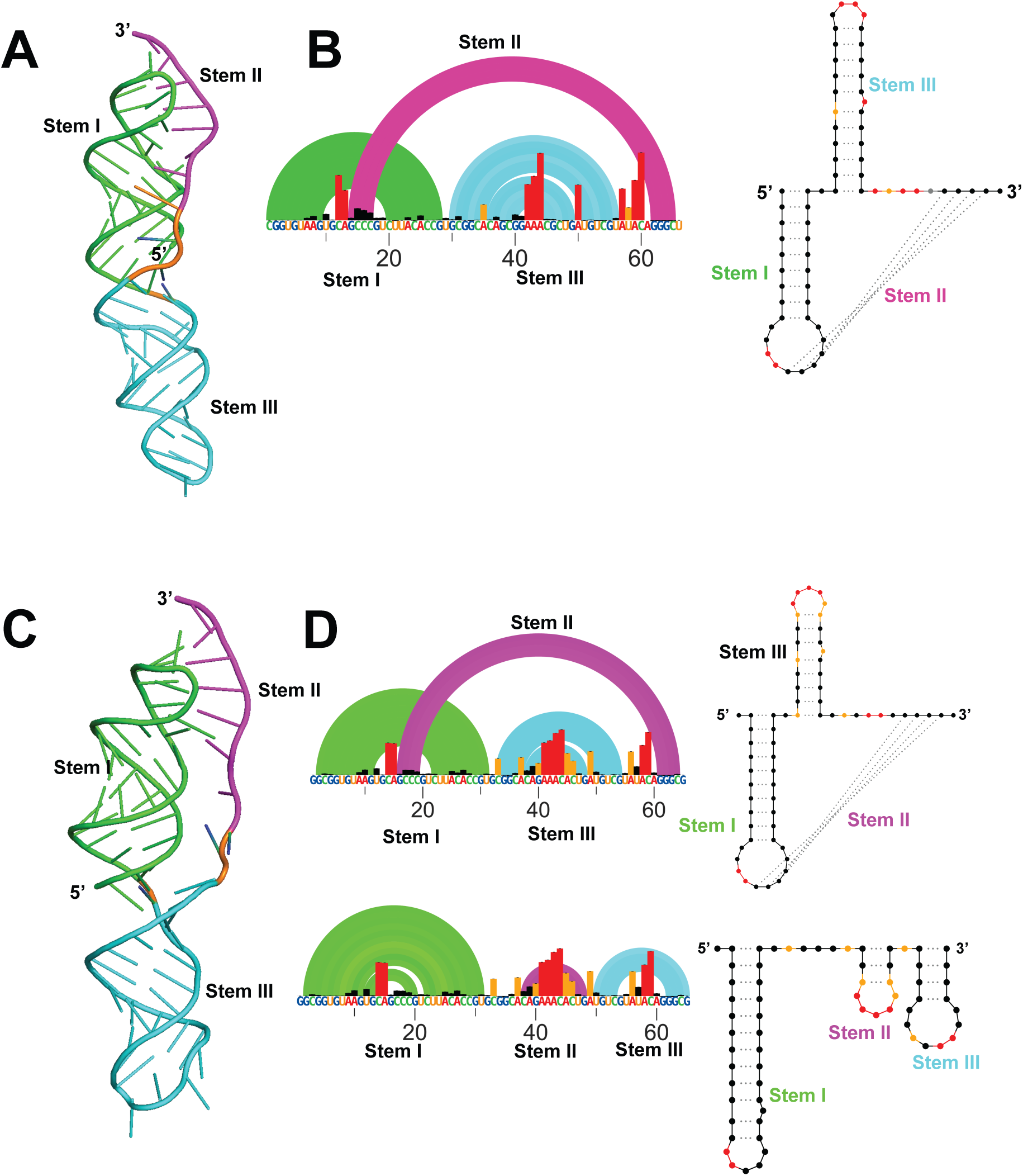
DMS-MaP for crystal structure constructs of SARS-CoV-2 FSE. (A) 3D crystal structure (PDB ID 7LYJ) of 66nt FSE construct in 3_6 H-type pseudoknot conformation (B) Arc plot and radial layout of 7LYJ computed through DMS-MaP and SHAPEknots in 3_6 topology (C) 3D crystal structure (PDB ID 7MLX) of 65nt FSE construct in 3_6 H-type pseudoknot conformation (D) Arc plot and radial layout of 7MLX computed through DMS-MaP and SHAPEknots in 3_6 topology. A minor 3_1 conformation for 7MLX is also present. Stem I, Stem II, and Stem III are annotated in the arc plot and radial layout.

### Predicted 77nt FSE conformation-stabilizing mutants uniformly decrease frameshifting efficiency

As mentioned above, our previous SHAPE chemical probe 5NIA (5-nitroisatoic anhydride) analysis on 77nt FSE (without the slippery sequence) also favors the 3_6 pseudoknot (Schlick, Zhu, Dey, et al. 2021; K. Zhang et al. 2021; Jones and Ferré-D’Amaré 2022; Roman et al. 2021). When alternatively probed by DMS, 3_6 emerges as the lowest energy conformation followed by 3_5 and 3_3 (Fig. S2, Table 1).

Using RAG-IF (inverse folding programs for graphs) (Fera et al. 2004), we have computationally designed three 77nt mutant systems with up to six mutations to selectively stabilize each of these three conformations. Our DMS data on these three constructs (Figures 3A, B and C) here confirm that each of these mutants selectively stabilize one of the three available conformations (Table 1). Indeed, SHAPEknots predicts a single conformation for each of these mutants (Table 1). Moreover, subjecting these 77nt mutant constructs to functional assays using dual luciferase reporter systems (see method section) reveals essentially abolished frameshifting (Figure 3D). This is surprising because despite the dominance of the 3_6 pseudoknot suggested by crystallography, Cryo-EM, SHAPE-MaP/DMS-MaP reactivity data, and computational predictions (Bhatt et al. 2021; Roman et al. 2021; K. Zhang et al. 2021; Schlick, Zhu, Dey, et al. 2021; Jones and Ferré-D’Amaré 2022), all mutants appear to lose frameshifting entirely. In fact, the abolished frameshifting from the 3_6 stabilizing mutant is the most puzzling result initially; however, this result can be explained by the required availability of other folds to the frameshifting process (see Discussion).

**Figure 3:**
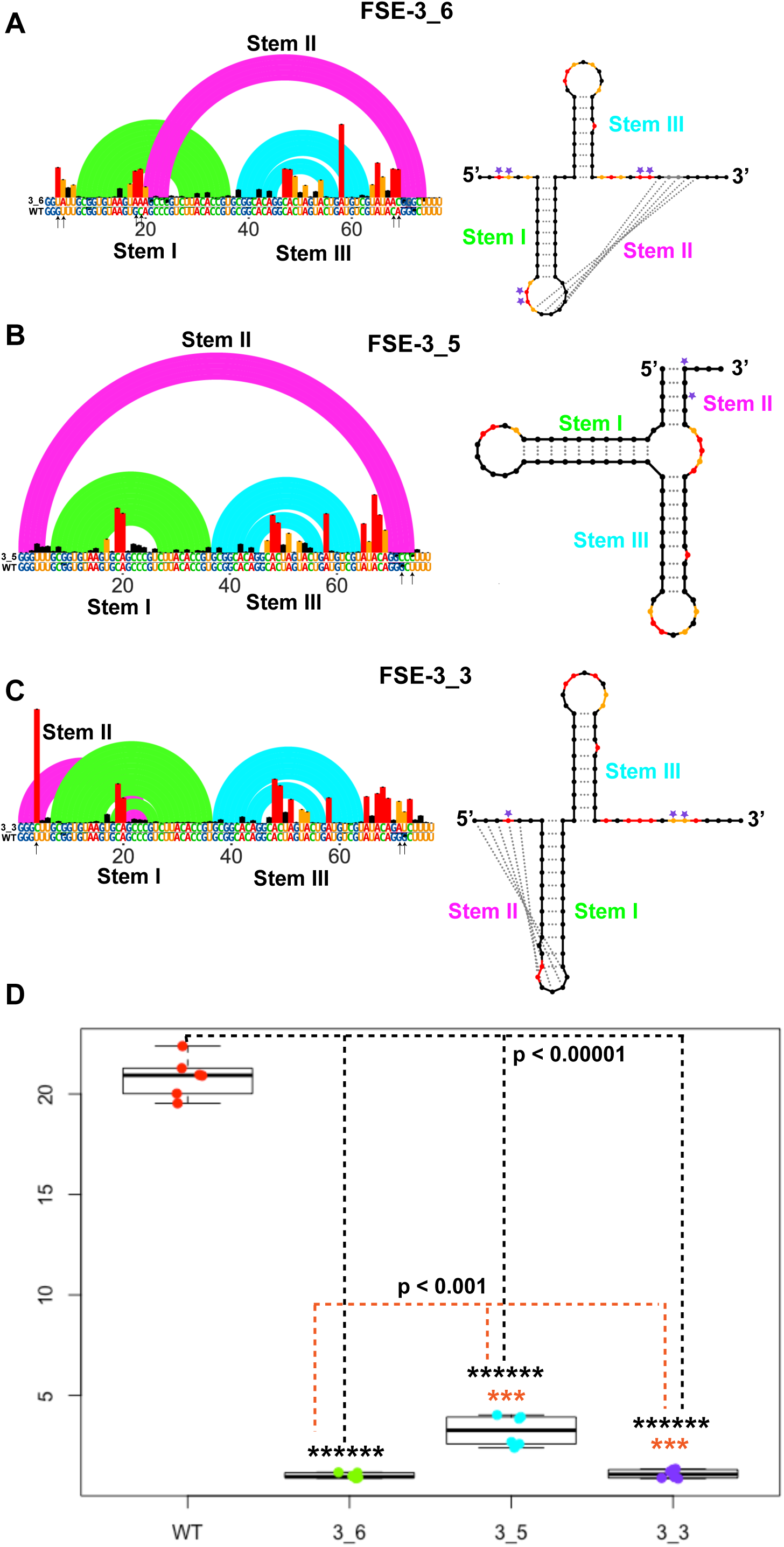
DMS-MaP and frameshifting efficiency of FSE conformation stabilizing mutants. Arc plot and radial layout computed by DMS-MaP and SHAPEknots for (A) FSE-3_6 mutants stabilizing 3_6 H-type pseudoknots (B) FSE-3_5 mutants stabilizing 3_5 three-way junction and (C) FSE-3_3 mutants stabilizing 3_3 H-type pseudoknots. Stem I, Stem II, and Stem III are annotated in the arc plot and radial layout. Arrows in arc plot and asterisk in radial layout represents point mutations. (D) frameshifting efficiency of 3_6, 3_5, and 3_3 stabilizing mutants highlighting substantial decrease in frameshifting efficiency when compared to WT 87nt FSE construct. Frameshifting efficiency was calculated using dual-luciferase assays.

As a control we also DMS probed, modeled, and measured the frameshifting efficiency of two additional longer WT constructs (156nt and 222nt long) (Figures 4A, B, and C, respectively). Our previous work has shown that longer constructs of SARS-CoV-2 FSE tend to adopt other conformations due to a competing Stem 1 and attenuator hairpin (Yan et al. 2022; Yan et al. 2023) (S. lee, S. Yan, A. Dey et al, Pers. comm., 2024). As can be seen in Figure 4C, both the 156nt and 222nt constructs have increased frameshifting compared to the short FSE. The 156nt construct show ∼15% more frameshifting efficiency while 222nt construct had ∼10% more frameshifting efficiency than 87nt WT construct (Fig. 4C). The 156nt construct, which according to DMS guided thermodynamic modeling adopts the three alternative conformations (Figure 4A, Table 2 and Table 4), has the highest frameshifting efficiency of all the constructs we tested. Even more puzzling, neither the 156nt nor 222nt constructs are predicted to adopt the 3_6 cryo-EM/crystal conformation (Table 2). The explanation here likely stems from the ease of refolding into related RNA folds during ribosomal translation (S. lee, S. Yan, A. Dey et al, Pers. comm., 2024, Figure 8); more in Discussion.

**Figure 4:**
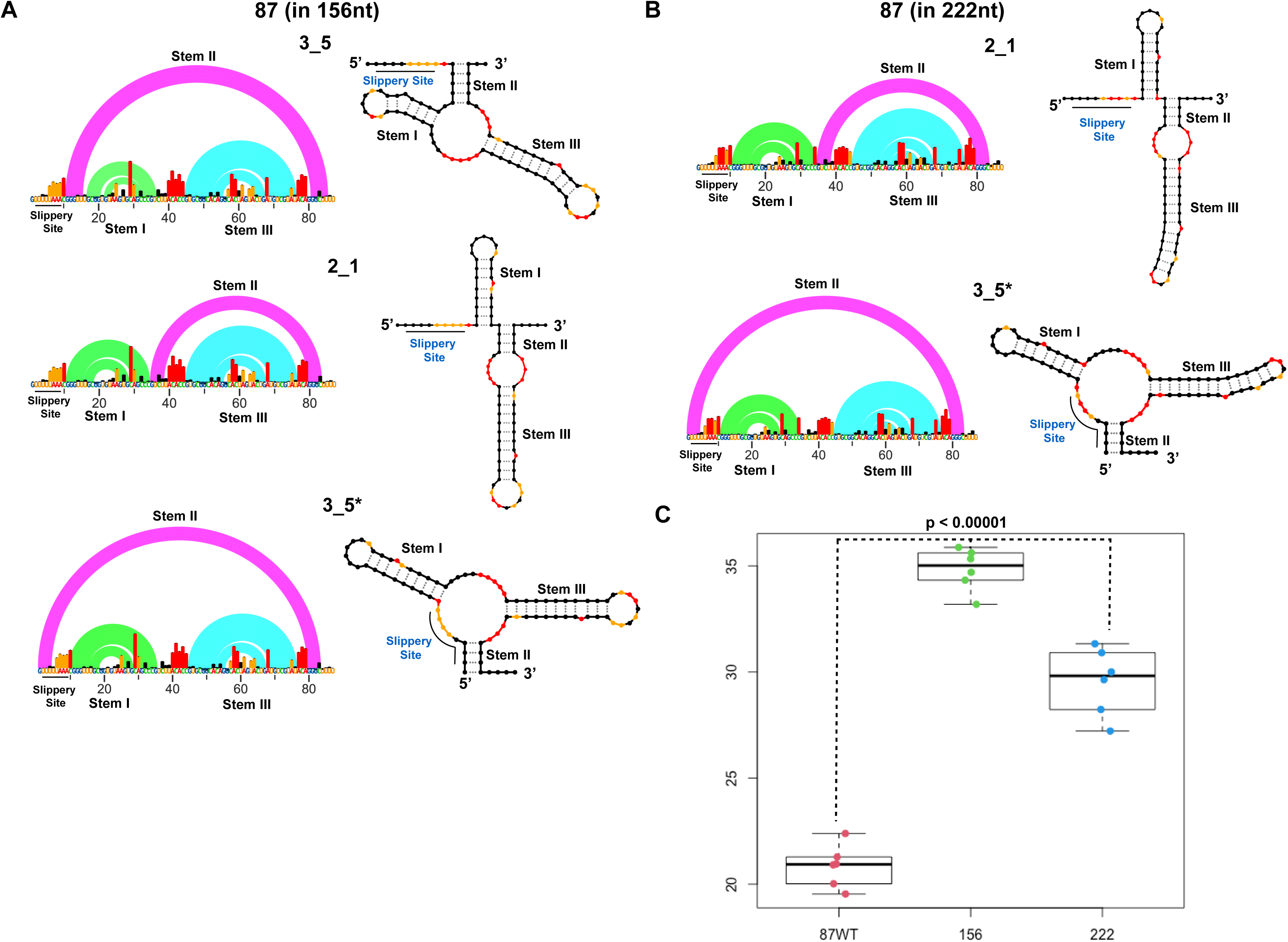
Conformational flexibility and frameshifting efficiency of FSE in larger constructs. Arc plot and radial layout computed by DMS-MaP and SHAPEknots for (A) 156nt construct in 3_5, 2_1, and 3_5* topology and for (B) 222nt construct in 2_1 and 3_5* topology. Slippery site, Stem I, Stem II, and Stem III are annotated in the arc plot and radial layout. (D) frameshifting efficiency of 156nt, and 222nt construct stabilizing mutants highlighting substantial increase in frameshifting efficiency when compared to WT 87nt FSE construct. Frameshifting efficiency was calculated using dual-luciferase assays.

**Table 2:**
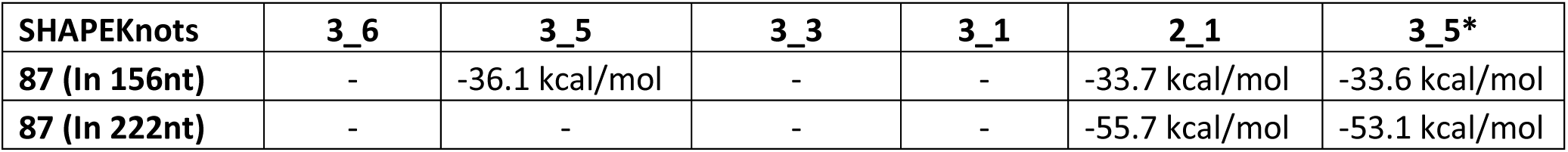
Minimum free energies computed for longer SARS-CoV-2 FSE constructs after DMS-MaP and ShapeKnots.

### Correlated mutational profiling for multiple conformations

So far, our analysis of FSE structures has relied on a traditional coupling of thermodynamic folding parameters with a pseudo-free energy term correction using DMS data. This approach is optimized for improved accuracy of minimum free energy predictions and was trained on highly structured RNAs such as the ribosomal RNAs (rRNAs). Yet, applications to model alternative secondary structures poses challenges, likely explaining the large differences in predicted free energies of folding across different constructs (Table 1 and Table 2). Three recently developed alternatives to thermodynamic structure modeling have been proposed to deconvolute alternative structures using correlated mutation in DMS-MaP data: DANCE-MaP (Olson et al. 2022), DREEM (Tomezsko et al. 2020), and DRACO (Morandi et al. 2021).

Analyses of our DMS-MaP data using these packages with SHAPEknots predictions yield conformer distributions shown in Tables 3 and 4. For the 87nt construct, DANCE-MaP, and DREEM computed 3_6 as predominant conformations (34%, and 89% respectively, Table 3). However, DRACO only predicted 16% of 3_6 conformation as minor conformation and 57% of 3_3 conformation as major fraction. In accordance with the above results DANCE- MAP also predominantly identified 3_6 conformation of 77nt construct (99%, Table 3). For 77nt FSE, all three programs identified 3_6 as the major conformation, but also suggest small (<6%) amounts of 3_5, 3_3 and 2_1 (2_1 is a simple two-stem structure).

**Table 3:** Boltzmann fractions computed for shorter SARS-CoV-2 FSE constructs using, Dance-MaP, DREEM, and DRACO.

|  |  | DANCE-MaP | DREEM | DRACO |
| --- | --- | --- | --- | --- |
| 87 (6XRZ) | 3_6 | 0.34 | 0.89 | 0.16 |
|  | 3_5 | 0.29 | 0.03 | 0.13 |
|  | 3_3 | 0.08 | 0.06 | 0.57 |
|  | 3_5* | 0.29 | 0 | 0 |
|  | 2_1 | 0 | 0 | 0.14 |
|  | 4_21 | 0 | 0.01 | 0 |
| 7LYJ (66nt) | 3_6 | 1 | 0.77 | 0.87 |
|  | 3_5 | 0 | 0 | 0 |
|  | 3_3 | 0 | 0 | 0 |
|  | 2_1 | 0 | 0.23 | 0.13 |
| 7MLX (65nt) | 3_6 | 1 | 0.78 | 0.68 |
|  | 3_5 | 0 | 0 | 0 |
|  | 3_3 | 0 | 0 | 0 |
|  | 3_1 | 0 | 0 | 0 |
|  | 2_1 | 0 | 0.22 | 0.19 |
|  | 2_2 | 0 | 0 | 0.13 |
| 77 | 3_6 | 0.99 | 0.935 | 0.657 |
|  | 3_5 | 0.007 | 0.0006 | 0.002 |
|  | 3_3 | 0.0004 | 0.057 | 0.145 |
|  | 2_1 | 0 | 0 | 0.195 |
| 3_6<br>Stablizing | 3_6 | 1 | 0.3 | 0.81 |
|  | 3_5 | 0 | 0 | 0.02 |
|  | 3_3 | 0 | 0 | 0 |
|  | 3_1 | 0 | 0.01 | 0 |
|  | 3_2 | 0 | 0 | 0.165 |
|  | 2_1 | 0 | 0.69 | 0 |
| 3_5<br>Stablizing | 3_6 | 0 | 0 | 0 |
|  | 3_5 | 1 | 1 | 0.9 |
|  | 3_3 | 0 | 0 | 0.03 |
|  | **3_1** | 0 | 0 | 0 |
|  | **3_2** | 0 | 0 | 0.07 |
| **3_3**<br>**Stablizing** | **3_6** | 0 | 0 | 0 |
|  | **3_5** | 0 | 0 | 0 |
|  | **3_3** | 1 | 0.8 | 0.01 |
|  | **3_1** | 0 | 0 | 0.004 |
|  | **3_2** | 0 | 0 | 0.05 |
|  | **2_1** | 0 | 0.2 | 0.27 |
|  | **3_8** | 0 | 0 | 0.67 |

For our structure stabilizing mutants, the combined analyses in Table 3 validate the changed distribution in the conformational landscape, except for the 3_6 mutant where only DREEM favors a simple two stem structure (2_1), and the 3_3 mutant, where only DRACO favors a 3_8 pseudoknot, which usually forms at long lengths due to competing alternative to stem 1 (S. lee, S. Yan, A. Dey et al, Pers. comm., 2024). Testing these three programs on the crystal structure data confirms these imperfect agreements.

Finally, our DMS-MaP data for longer constructs using these packages suggest a variety of conformations, including 3_6, 3_5, 2_1, 3_3, and 3_2 (Table 4). These also includes some additional conformational variants like 3_5*, 2_1*, and 3_2* which are similar to 3_5, 2_1, and 3_2 topology with only differences in base pairing arrangements (Table 4). This increase of folds at larger lengths is consistent with our recent finding of structural heterogeneity at increasing FSE lengths (S. lee, S. Yan, A. Dey et al, Pers. comm., 2024).

**Table 4:**
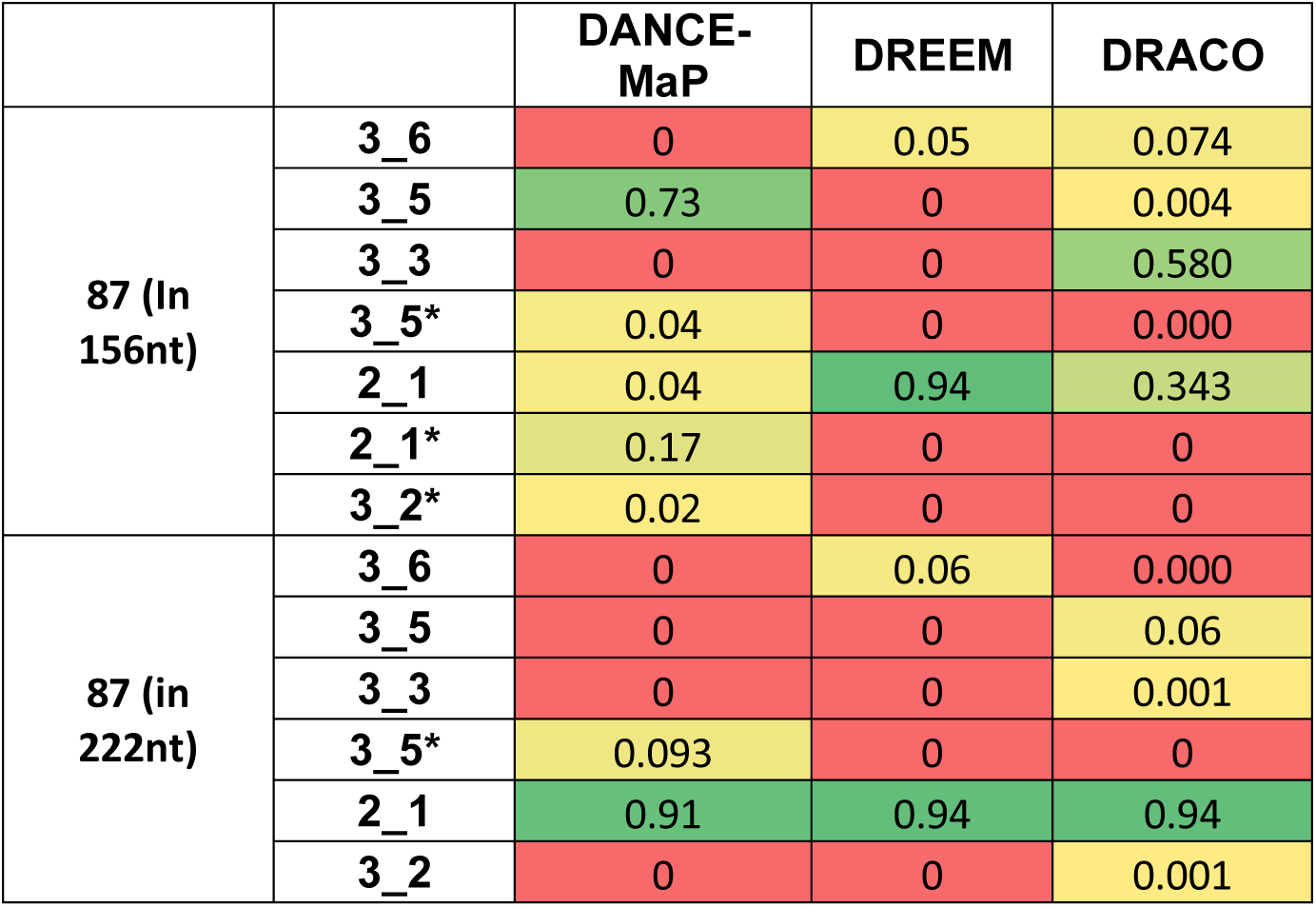
Boltzmann fractions computed for longer SARS-CoV-2 FSE constructs using, Dance-MaP, DREEM, and DRACO.

## Discussion

As discussed in this work, the SARS-CoV-2 FSE not only plays a critical role in coronavirus replication and virulence (K. Zhang et al. 2021; Sanders et al. 2020; Iserman et al. 2020; Schlick, Zhu, Jain, et al. 2021; Schlick, Zhu, Dey, et al. 2021; Yan et al. 2022; Yan et al. 2023; Jones and Ferré-D’Amaré 2022; Bhatt et al. 2021; Roman et al. 2021), but is also of intense interest because of the intriguing repertoire of length dependent conformations. Prior studies have reported many possible conformations (K. Zhang et al. 2021; Bhatt et al. 2021; Jones and Ferré-D’Amaré 2022; Roman et al. 2021; Manfredonia et al. 2020; Iserman et al. 2020; Sanders et al. 2020; Schlick, Zhu, Dey, et al. 2021) for the SARS-CoV-2 FSE, although cryo-EM and crystallographic studies have identified the H-type pseudoknot (3_6) at short lengths (K. Zhang et al. 2021; Jones and Ferré-D’Amaré 2022; Roman et al. 2021). Our work here has probed the FSE structural abundance by subjecting several wildtype sequences (77nt, 87nt, 156nt and 222nt) and three predicted structure-stabilizing mutants (Schlick, Zhu, Dey, et al. 2021) to dimethyl sulfate probing and mutational analysis profiling (DMS-MaP). By using these experimental data and analysis packages designed to handle multiple conformations (DANCE-MaP (Olson et al. 2022), DREEM (Tomezsko et al. 2020), and DRACO (Morandi et al. 2021)), we characterized the structural distributions in these variable systems. We also performed functional assays for the mutants and showed abolished frameshifting. The wildtype system characterization emphasized the prevalence of the 3_6 pseudoknot but also the emergence of alternative forms: 3_3 H-type pseudoknot and 3_5 three-way junction (Fig. 1, 2 and S2, Table 1 and Table 3).

These findings are in agreement with our earlier SHAPE studies (Schlick, Zhu, Dey, et al. 2021) and a recent single-molecule structural study where the architecture of core FSE (86nt) element was divided in two cluster conformations with 3_6 and 3_5 as dominant topology (Pekarek et al. 2023). These findings also agree with our recent characterization of length-dependent FSE systems that show the emergence of alternative stem 1 along with simple stem loop folds (3_1, 3_2, 2_1) at long lengths (S. lee, S. Yan, A. Dey et al, Pers. comm., 2024). Clearly SARS-CoV-2 FSE is highly flexible in nature and can exist in multiple conformations.

The extension of these analyses to the three RAG-designed mutants (Schlick, Zhu, Dey, et al. 2021) suggests markedly different distributions, namely, emphasizing the dominance of the 3_6, 3_3, and 3_5 conformations in turn, although the three deconvolution analysis packages show variations. Intriguingly, the essential loss of frameshifting efficiency for these mutants (Fig. 3D) suggests that suppression of structural transitions, or the ability of the RNA to sample alternative conformations, is essential for frameshifting. Indeed, our conformational landscape viewpoint has revealed essential residues to stabilize central conformations while others to add variability into the FSE conformational repertoire (Yan et al. 2023). But while it is plausible that 3_5 and 3_3 stabilizing mutants may not frameshift, why would the 3_6 stabilizing mutant does not frameshift significantly?

The frameshifting ability must be an intricate function of the folding and refolding of variable FSE RNA segments interacting with multiple ribosomal RNA during translation. As we proposed in (S. lee, S. Yan, A. Dey et al, Pers. comm., 2024), the 3_6 conformation is dominant at short lengths when the slippery site is constrained and the 5’-end sequence from FSE is short. Otherwise, alternative stem 1 and attenuator hairpin AH consort to block the pseudoknot 3_6 and favor 2_1 or 2_2 stem loops and/or a different pseudoknot 3_8 (S. lee, S. Yan, A. Dey et al, Pers. comm., 2024). Once the 3_6 pseudoknot is recovered upon ribosomal unwinding, it competes with the 3_3 pseudoknot through structural interconversions (Yan and Schlick, in preparation). All these structural interchanges must occur fluidly and quickly, involving energy barriers that are lowered by ribosomal intervention and winding/unwinding forces. Thus, a new intervention avenue for suppressing infection might involve tampering with this conformational cascade via mutations and/or structure stabilizing small drug compounds. Pursuing such designs by experiments and computations using FSE systems that yield specific conformational distribution will be fascinating.

Finally, the notion that RNAs are versatile molecules that adopt many possible conformations has been well appreciated for riboswitches and non-coding long RNAs (Bose, Saleem, and Mustoe 2024; Serganov and Patel 2012). Yet characterizing their conformational variability is challenging. DMS-data deconvoluting packages as used here certainly help examine these possibilities, but results among programs are not consistent, pointing to many areas of future improvement. Clearly, considering an extended landscape of length-dependent structures with various programs along with other experimental and computational data is important. Viruses like coronaviruses or HIV (Mouzakis et al. 2013) provide particularly interesting examples of the variability of structures found in very short segments. Indeed, that both uses the -1 frameshifting mechanism suggests that frameshifting viral systems define ideal platforms for interrogating the intricate landscape of multiple RNA folds. Our study has thus emphasized the importance of these multiple forms and the challenges associated with their analyses. New therapeutic strategies to affect frameshifting also naturally emerge.

## Materials and Methods

### SARS-CoV-2 sequence

29891nt SARS-CoV-2 reference sequence was retrieved from GISAID (Elbe and Buckland-Merrett 2017) (Accession ID: EPL_ISL_402124). The 87nt FSE occupies the residues 13462-13548.

### *In vitro* RNA chemical probing read by mutational profiling

Various SARS-CoV-2 FSE constructs were synthesized as G-blocks from Integrated DNA Technologies (IDT). These constructs were flanked at both 5’ and 3’ end by RNA cassettes (Wilkinson, Merino, and Weeks 2006). T7 promoter region was added to the 5’ end of each construct for *in vitro* transcription of RNA using T7 RNA polymerase from T7 HiScribe RNA synthesis kit (New England Biolabs). Synthesized RNA was subjected to DNase treatment (TURBODNase) and was further purified using Purelink RNA mini kit (Invitrogen) and quantified using nanodrop.

For chemical probing, 6 μg of purified *in vitro* transcribed RNA was denatured at 65 °C for 5 min and snap-cooled in ice. Following denaturation, folding buffer (100 mM KCl, 10 mM MgCl2, 100 mM Bicine, pH 8.3) was added to the denatured RNA and the whole reaction was incubated at 37 °C for 10 min. The folded RNA was further treated with 10 μl of 1:10 ethanol diluted Dimethyl Sulfate (DMS). For control, equivalent volume of ethanol was added to the folded RNA. Probing was initiated by incubating the reaction mixture at 37 °C for 5 min and was quenched afterwards by adding 100 μl of 20% β-mercaptoethanol (β-ME). Modified and un-modified RNAs were purified using Purelink RNA mini kit and quantified using nanodrop.

### Library construction, sequencing, and data processing

Following chemical probing, both modified and un-modified RNA were reverse transcribed using specific primer complementary to the 3’ RNA cassette (Dey 2023) and Superscript II reverse transcriptase under error prone conditions as previously described (Smola et al. 2015). The generated cDNA was purified using G50 column (GE healthcare) and subjected to second strand synthesis (NEBNext Second Strand Synthesis Module). For constructing next-generation sequencing libraries, the double-stranded (ds) cDNA was PCR amplified with primers directed against 5’ and 3’ RNA cassettes and NEB Q5 HotStart polymerase (NEB). To introduce unique barcodes secondary PCR was performed using TrueSeq primers (NEB) (Smola et al. 2015). Amplified products were purified using Ampure XP (Beckman Coulter) beads and quantification of the libraries were done using Qubit dsDNA HS Assay kit (ThermoFisher). Purified libraries were quality checked using Agilent Bioanalyzer. These libraries were finally sequenced as 2 x 151 paired end read on Illumina MiSeq platform.

Shapemapper2 algorithm (Busan and Weeks 2018) was used to calculate mutation frequency in both chemically modified (DMS treated) and control/un- modified (ethanol treated) RNA samples. Chemical modifications on each RNA nucleotide was calculated using the following equation:

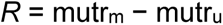

where R is the chemical reactivity, mutr_m_ is the mutation rate calculated for chemically modified RNA and mutr_u_ is the mutation rate calculated for un- modified control RNA samples (Smola et al. 2015; Smola and Weeks 2018). Shapemapper2 was also used to calculate the parse mutations from the sequencing data. Chemical reactivity obtained from Shapemapper 2.0 was used to inform a minimum free energy structure using ShapeKnots (Hajdin et al. 2013). Resultant model was visualized using VARNA (Darty, Denise, and Ponty 2009). RNA arc and secondary structure models were generated using RNAvigate (Irving and Weeks 2023).

### DANCE-MaP analysis on SARS-CoV-2 FSE constructs

DANCE-MaP (deconvolution and annotation of ribonucleic conformational ensembles) algorithm uses maximum likelihood (ML) strategy to extract large amount of total information available from single chemical probing experiment (Olson et al. 2022). DANCE-MaP, measures chemical reactivities on all four nucleotides simultaneously including base pairing and tertiary interactions which enables it to measure and detect possible RNA ensembles at single nucleotide resolution. Parsed files (for both modified and un-modified datasets) were used in DANCE-MaP pipeline to calculate and validate alternative conformations for SARS-CoV-2 FSE constructs. The resultant reactivity profile was then used as a constraints in ShapeKnots (Hajdin et al. 2013), to generate probable SARS-CoV- 2 FSE ensembles for different constructs.

### DREEM analysis on SARS-CoV-2 FSE constructs

”Detection of RNA folding ensembles using expectation–maximization” (DREEM) algorithm (Tomezsko et al. 2020) was run directly on the sequencing reads of DMS modified SARS-CoV-2 FSE constructs to determine the presence of alternative structures in the constructs. Using an expectation-maximization technique, DREEM clusters the sequencing reads into discrete groups based on patterns of DMS-induced mutations. Log-likelihood is maximized to obtain the DMS modification rate per base for each cluster. In this study, a maximum of K=3 clusters were used to group the bit-vectors. The resulting DMS reactivities for each cluster were then used as constraints for ShapeKnots (Hajdin et al. 2013) predictions. Hence, distinct structural clusters with their relative ratios result in different folds, which represents the heterogeneity of RNA secondary structure.

In order to predict the 87nt FSE structure for long constructs of 156 and 222nt, DMS reactivity constraints of the 87nt region retrieved from the normalized reactivity profile by DREEM for the complete constructs were used.

### DRACO analysis on SARS-CoV-2 FSE constructs

We also applied the DRACO algorithm (Morandi et al. 2021), which performs deconvolution of alternative RNA conformations from mutational profiling experiments with a combination of spectral clustering and fuzzy clustering, to validate the structure predictions of SARS-CoV-2 FSE. Spectral clustering is performed for the sliding windows along the transcript, allowing the optimal number of coexisting conformations (clusters) to be automatically identified from the eigen gaps. Following the determination of the number of clusters, fuzzy clustering is carried out to allow bases to be weighted according to their affinity to each cluster. DRACO then reconstructs overall mutational profiles by merging overlapping windows with the same number of clusters. DRACO reports consecutive sets of windows with varying amounts of clusters separately. The pair-end reads were merged by pear (J. Zhang et al. 2014) and mapped to the reference sequence using the rf-map tool (Incarnato et al. 2018) (parameters: -b2 -cqo -ctn -mp ’–very-sensitive local’). Resulting BAM files were then analyzed with the rf-count tool to produce MM files (-r -m -mm -na -ni). MM files were analyzed with DRACO (Morandi et al. 2021) (parameters: – allNonInformativeToOne –nonInformativeToSurround –minClusterFraction 0.1) and deconvoluted mutation profiles were extracted from the resulting JSON files. Normalized reactivity profiles were obtained by first calculating the raw reactivity scores via the scheme by DMS-MaP (Zubradt et al. 2017) as the per-base ratio of the mutation count and the read coverage at each position, and then by 90% Winsorizing as normalization method, using the rf-norm tool (Incarnato et al. 2018) (parameters: -sm 4 -nm 2 -rb AC -mm 1). Data-driven RNA structure prediction was performed using ShapeKnots (Hajdin et al. 2013) and the normalized reactivity profiles.

### Dual Luciferase Assay of -1 PRF

pJD2514-87, a test SARS-CoV-2 reporter plasmid of 87nt construct and pJD2257 a 0-frame readthrough control plasmid for dual luciferase reporter assay was obtained as a kind gift from Prof. Jonathan D. Dinman, University of Maryland. Q5 site directed mutagenesis kit (NEB) was used to generate various SARS-CoV-2 FSE conformation stabilizing constructs and other length dependent constructs and their sequences were confirmed through sanger sequencing (Eurofins genomics). HEK293 cells were used as host cells to transfect the SARS-CoV-2 FSE dual luciferase reporter plasmid using Lipofectamine 2000 transfection reagent (ThermoFisher Scientific). The -1 PRF efficiency was assayed in cultured HEK293 as described previously (Kelly et al. 2020) using dual-luciferase reporter assay system kit (Promega). 24 hr post transfection, HEK293 cells were washed with 1X PBS buffer and then lysed with 1X passive lysis buffer (Promega). The cells were cleared of any cell debris by centrifugation at 14000 rpm for 10 min at 4°C. Each cell lysate both test [pJD2514] and 0-frame read through control [pJD2257] were assayed in triplicate in 96-well plate, and luciferase activity was quantified using CLARIOstar (BMG Labtech) plate reader. SARS-CoV-2 frameshifting efficiency was calculated by averaging the triplicates of firefly and *Rennila* luciferase per samples and calculating the ratio of firefly to *Rennila* luciferase of each sample. Percent frameshifting was subsequently calculated for test sample by comparing their luminescence ratio (as calculated above) with the 0-frame readthrough control set at 100%. This ratio of ratios provided the percent -1 PRF efficiency of each SARS-CoV-2 FSE constructs. Statistical analyses were conducted through Welch two sample T-test using R software.

## Supporting information

SARSCOV2_FSE_SNRSAM file

## Acknowledgments

This work was supported by U.S. National Institutes of Health grants NHLBI R01 HL111527 and NIGMS R35 GM140844 to A.L. and National Science Foundation Award DMS -2151777 from the Division of Mathematical Sciences to T.S. and A.L, National Institutes of Health R35GM122562 from the National Institute of General Medical Sciences to T.S. Dr. Abhishek Dey acknowledges the Department of Biotechnology, Govt. of India for the Ramalingaswami Re-entry fellowship (BT/RLF/Re-entry/02/2021). The authors also acknowledge Prof. Jonathan D. Dinman for kindly providing reporter plasmids for dual luciferase assays to assess -1 PRF efficiency of SARS-CoV-2 FSE constructs.

## Supporting Information

DMS reactivities calculated for all the SARS-CoV-2 FSE constructs are available as SARSCoV-2-FSE_SNRSAM.xlsx. SRA metadata for all the constructs (BioProject ID: PRJNA1086021) can be accessed from NCBI using the following link: http://www.ncbi.nlm.nih.gov/bioproject/1086021.

---

**Figure S1:** Statistical analysis between replicates of various SARS-CoV-2 FSE constructs. Scatter plot with Pearson correlation coefficient showing higher reproducibility between the replicates for (A) 77nt (B) 3_6 stabilizing mutant (C) 3_5 stabilizing mutant (D) 3_3 stabilizing mutant (E) 87nt (F) 156nt and (G) 222nt FSE constructs.

**Figure S2:** Conformational flexibility in the 77nt SARS-CoV-2 FSE construct. Arc plot and radial layout computed by DMS-MaP and SHAPEknots for 77nt FSE construct depicting (A) 3_6 H-type pseudoknot (B) 3_5 three-way junction (C) 3_3 H-type pseudoknot. Stem I, Stem II, and Stem III are annotated in the arc plot and radial layout.

## Notes

http://www.ncbi.nlm.nih.gov/bioproject/1086021.

### Competing Interest Statement

The authors have declared no competing interest.

