## Supplementary figures and images for "Abolished frameshifting for predicted structure-stabilizing SARS-CoV-2 mutants: Implications to alternative conformations and their statistical structural analyses"

### Supplementary Figure 1

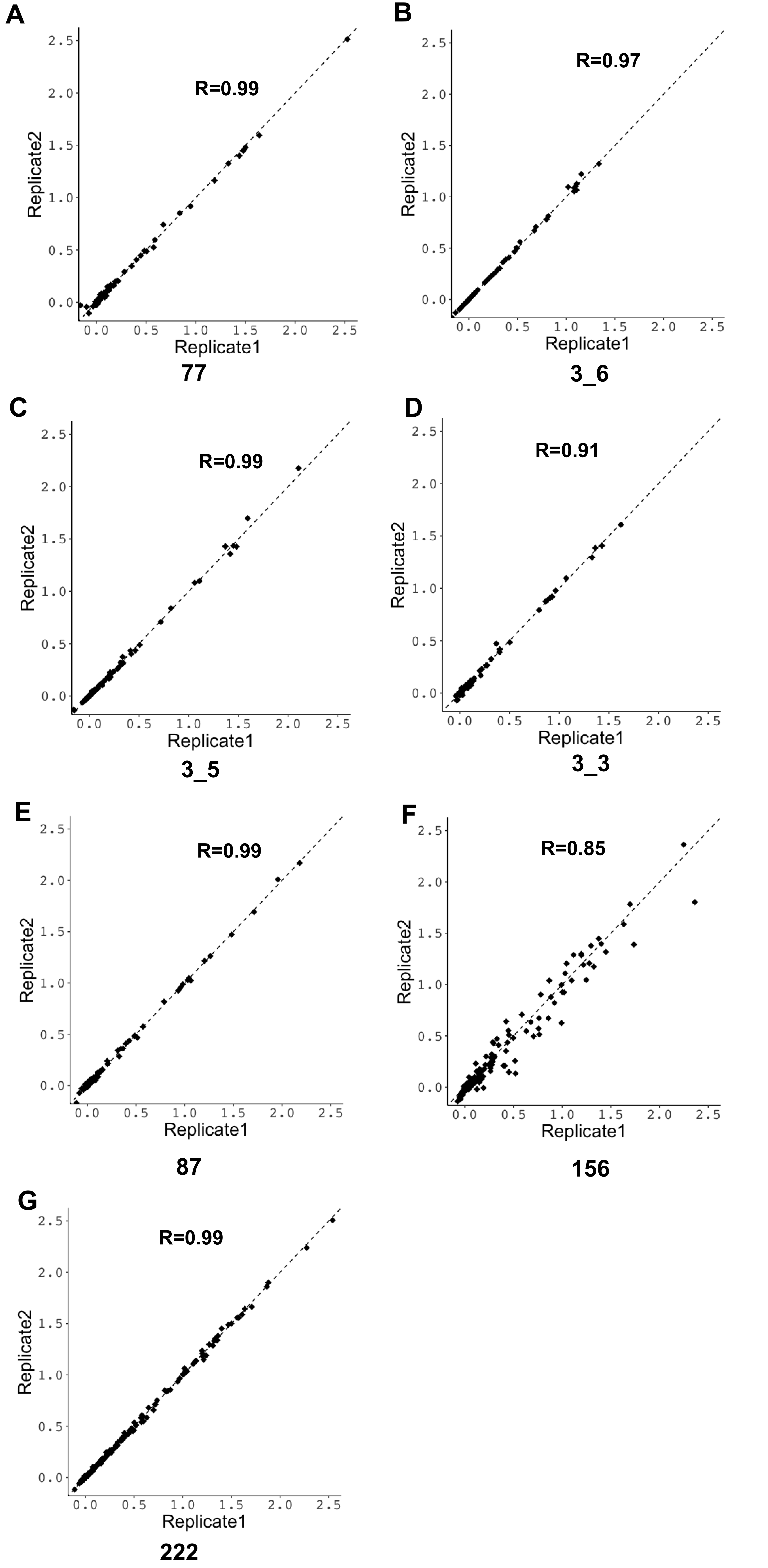

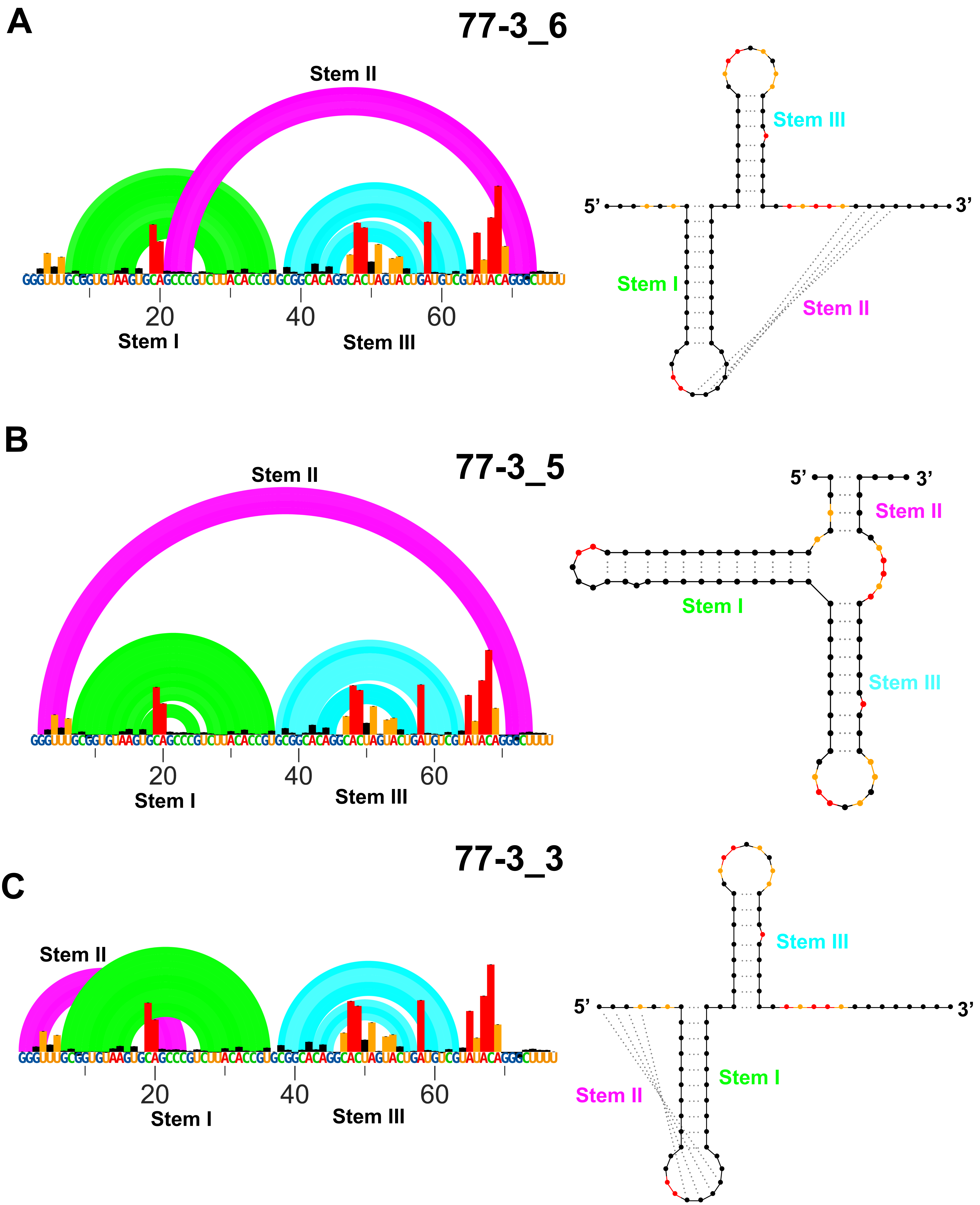

### Supplementary Figure 2

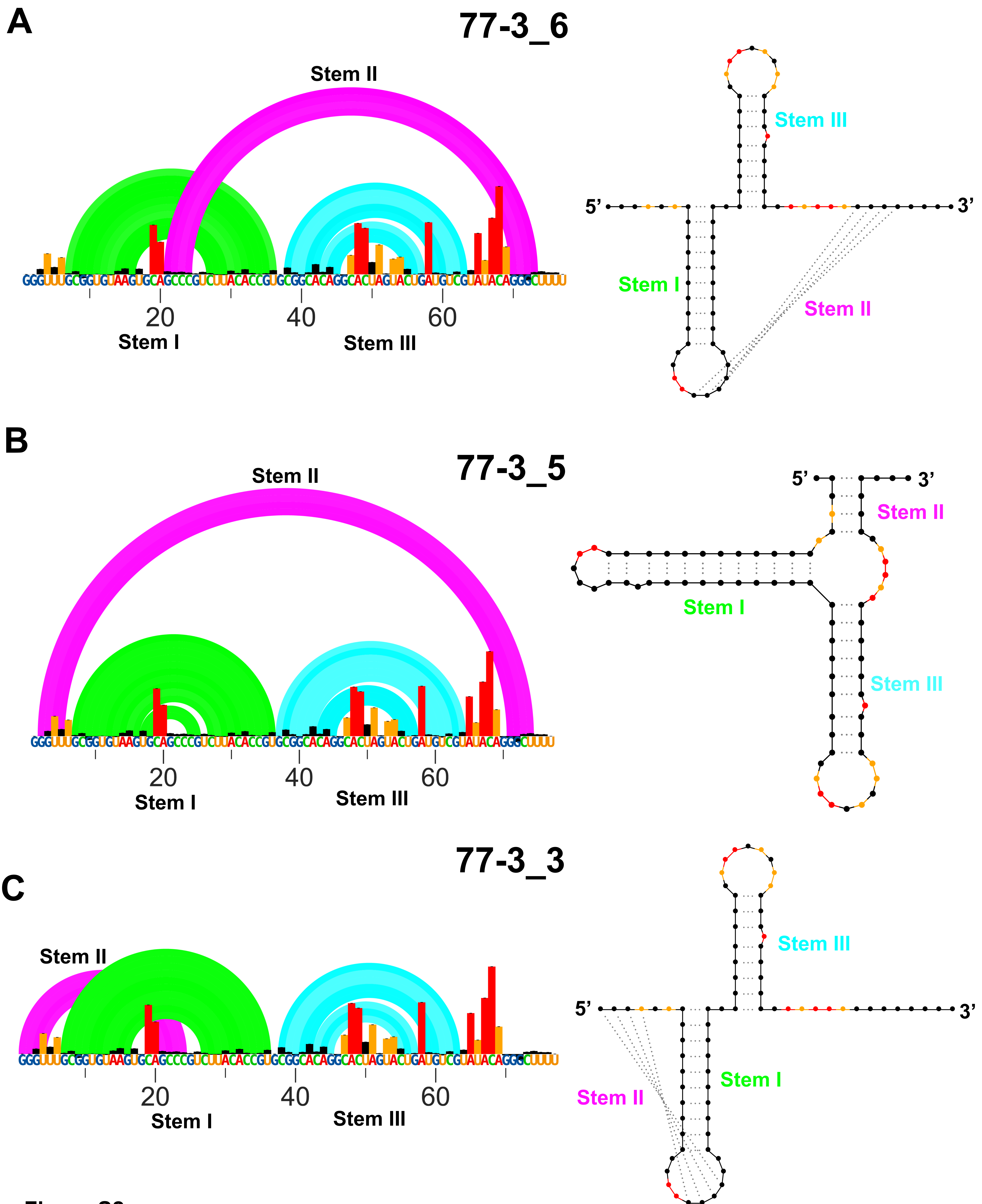
